# Nanoparticle targeting of mechanically modulated glycocalyx

**DOI:** 10.1101/2023.02.27.529887

**Authors:** Afia Ibnat Kohon, Kun Man, Katelyn Mathis, Jade Webb, Yong Yang, Brian Meckes

## Abstract

The mechanical properties and forces in the extracellular environment surrounding alveolar epithelial cells have the potential to modulate their behavior. Particularly, breathing applies 3-dimensional cyclic stretches to the cells, while the stiffness of the interstitium changes in disease states, such as fibrosis and cancer. A platform was developed that effectively imitates the active forces in the alveolus, while allowing one to control the interstitium matrix stiffnesses to mimic fibrotic lung tumor microenvironments. Alveolar epithelial cancer cells were cultured on these platforms and changes in the glycocalyx expression were evaluated. A complex combination of stiffness and dynamic forces altered heparan sulfate and chondroitin sulfate proteoglycan expressions. Consequently, we designed liposomal nanoparticles (LNPs) modified with peptides that can target heparan sulphate and chondroitin sulfates of cell surface glycocalyx. Cellular uptake of these modified nanoparticles increased in stiffer conditions depending on the stretch state. Namely, chondroitin sulfate A targeting improved uptake efficiency in cells experiencing dynamic stretches, while cells seeded on static stiff interstitium preferentially took up heparan sulfate targeting LNPs. These results demonstrate the critical role that mechanical stiffness and stretching play in the alveolus and the importance of including these properties in nanotherapeutic design for cancer treatment.

## Introduction

Lung cancer is the leading cause of cancer-related death, worldwide with mortality rates of 22% in 2022.^1^ High mortality rates stem from a combination of limited screening,^2^ highly metastatic potential of cells,^3,4^ relapse after common chemotherapeutic treatments,^5^ and absence of efficient alternative treatment options. In response, nanotherapeutics are being developed to improve drug accumulation in tumor sites while reducing off-target effects.6–10 Nanoparticles as therapeutic carriers for lung cancer are appealing because their unique size and structure designs allow them to maintain stability, cross biological barriers,^11^ bind to multiple targeting ligands,^12^ reduce non targeting cytotoxic effects of drugs,^13^ and improve circulation times with improved loading capacity^14^ and releasing efficiency.^15–18^ Although, targeted delivery of modified nanocarriers to lung cells has been developed with the purpose of improving therapeutic outcomes,^19–23^ few have reached clinical relevance^24^ due in part to the unique delivery challenge that is imparted by the physical environment of the lung and absence of efficient alternative options.

The structure and mechanics of the extracellular matrix (ECM) plays a vital role in many cellular physiological activities, such as cellular growth, proliferation, migration, and differentiation.^25–27^ In the lung alveolus, alveolar epithelial cells are exposed to 3-dimensional (3-D) stretching cycles that coincide with breathing. Conventional culture models used to study nanotherapeutic delivery to alveolar cells do not effectively mimic this dynamic environment, thus they fail to reflect mechano-transmitted cues that play essential roles in many disease etiologies.^28– 31^ In lung cancer, the tumor microenvironment is remodeled into a fibrotic phenotype.^32^ Specifically, the enhanced action of fibroblasts, fibrocytes,^33–36^ and mesenchymal stems cells^37^ in tumor microenvironment causes stiffening of the interstitial ECM,^38,39^ on which epithelial cells reside. This stiffening process is known to affect how nanomaterials interact with epithelial cells and lung fibroblasts as increased stiffness promotes uptake of Au nanoparticles (NPs) while reducing uptake of carbon nanotubes.^40–44^ However, designs that take advantage of substrate driven differences in uptake have not been realized nor have the effects of dynamic forces been included.

Recent studies have highlighted the connection between tumor mechanical environment and the cell surface glycocalyx^45–48^ – a dense sugar layer on the surface of cell membranes that acts as a mediator between cells and their environment. Changes in glycocalyx density and gene expressions were observed in some cancer cells on stiffer substrates. In the lung epithelium, the glycocalyx is composed primarily of hyaluronic acid, heparan sulfates, and chondroitin sulfates.^49,50^ However, little is known about stiffness related changes in lung cancer epithelium glycocalyx and how this can be leveraged for nanoparticle targeting.

To assess how fibrotic lung interstitium changes glycocalyx structures and nanoparticle entry, we adapted a recently reported cell culture platform^51^ that applies a physiological 3-D mechanical stretch to lung cancer epithelium. Cells grown on these platforms better mimic lung tissue properties such as tight junction formation, incident forces, and intercellular and cell-substrate interactions of lung epithelial cells. Using the biomimetic devices, we show that dynamic stretch forces combined with stiffness changes in the interstitium alter glycocalyx gene expression. In response, we synthesized lipid nanoparticles with specific glycocalyx recognition peptides on the surfaces. Specifically, chondroitin sulfate A (CSA) targeting liposomes imposed better uptake efficiency in a fibrotic, mechanically stretched environment of cancerous lung cells. In contrast, heparan sulphate targeting improved uptake in static fibrotic conditions. Together, these results highlight the importance of biomimetic dynamic stretch in designing nanoparticle therapeutics.

## Results & Discussion

### Fabrication and Characterization of Dynamic Alveolar Interstitium Platform

In order to understand how mechanical environment and dynamic forces impact cell interactions with nanoparticles, we created a dynamic platform consisting of a cell culture chamber, an interstitium layer, and a pneumatic chamber (**Figure 1**), made of polydimethylsiloxane (PDMS). A biomimetic nanofibrous membrane was placed on top of the interstitium layer, which was sandwiched between the cell culture chamber and the pneumatic chamber (**Figure 1B-E**). The assembly between these parts was completed by using a PDMS thin film adhesive as we previously reported^52^ (See the Methods).

**Figure 1.**
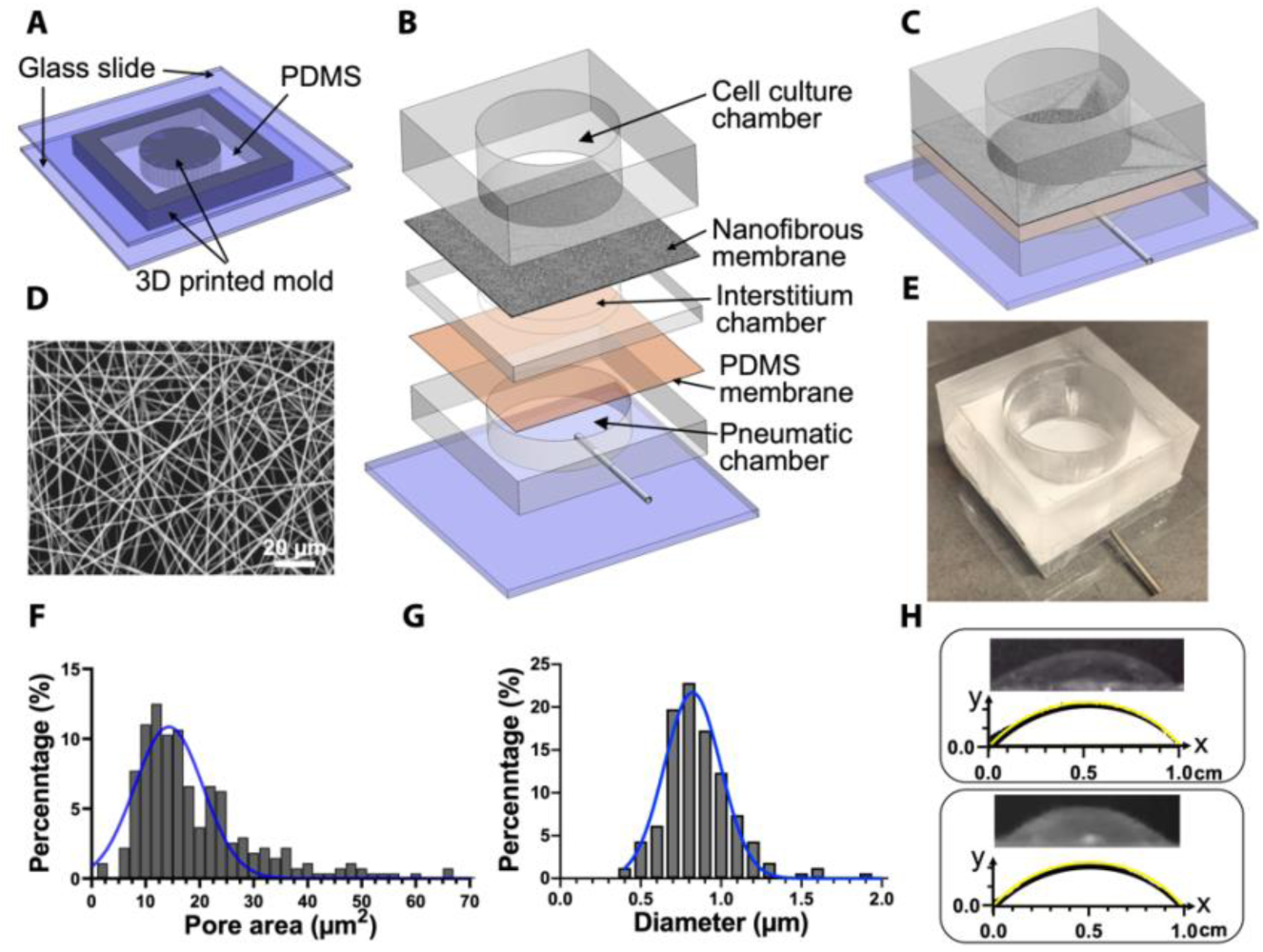
Design and characterization of the platform. **A)** Fabrication of the layer for cell culture, interstitium, and pneumatic chambers. The structures of the layers were the same with different heights (8mm for cell culture layer, 1mm for interstitium layer and 4mm for pneumatic layer). **B)** Exploded illustration of the platform fabrication. **C)** Illustration of the assembled platform. **D**) SEM image of the nanofibrous membrane. **E**) Image of the platform. **F**) Pore size distribution and **G)** fiber diameter distribution of the nanofibrous membrane. **H)** Characterization of the 3-D mechanical stretch. Original CCD images and the profiles (black curve) of the deformed PDMS membrane bonded on the pneumatic chamber (upper frame) or nanofibrous membrane on the interstitium bonded on the pneumatic chamber (lower frame) under a theoretical strain of 15% (yellow curve). The scales of axis x and y were the same.

The nanofibrous membrane was generated by electrospinning a polycaprolactone (PCL) solution to mimic the nanofibrous basement membrane of alveolar epithelium since PCL nanofibrous membrane has been reported to support epithelium formation and favor cell-cell crosstalk.^53–55^ The nanofibrous membrane had an average diameter of 820 ± 180 nm and an average pore size of 14.29 ± 6.33 μm^2^ (**Figure 1D,F-G**). The interstitium layer was prepared with PDMS and its stiffness was determined by tailoring the type and composition of PDMS to replicate the stiffness of normal (1-5 kPa) and fibrotic (20-100 kPa) lung tissues,^56–58^ *i*.*e*., PDMS Sylgard 527 for 5 kPa and a 10:1 (w/w) mixture of Sylgard 527 and 184 for 50 kPa, respectively.^59^ The nanofibrous membrane was first oxygen plasma treated and coated with type I collagen, and then immobilized on the PDMS interstitium, thus allowing the cells to adhere to the nanofibers but not onto the PDMS upon seeding. As such, the nanofibers provided contact guidance for the cell to adhere and grow, while the interstitium only regulates the cells with its stiffness. Moreover, to imitate the breath movement, a theoretical 15% linear strain was applied at a frequency of 0.2 Hz. When air was infused to the pneumatic chamber, the experimental observations of the strains of the PDMS membrane and the nanofibrous membrane immobilized on the interstitium agreed with the theoretical calculation (**Figure 1H**), indicating that the 3-D mechanical stretch was transmitted through the interstitium to the nanofibrous membrane, which provided mechanical stimulation to the adherent cells. Taken together, this platform provides a biomimetic system for evaluating nanoparticle targeting.

### Lung Epithelium Glycocalyx Gene Expression Changes in Response to Mechanical Environment

With the goal of designing nanoparticles that target cells based partially on the cellular response to mechanical cues, we assessed changes in surface related proteins. Since lung epithelial glycocalyx changes are linked to cancer and fibrosis,^38,39,60^ conditions where the mechanical properties of the interstitium are altered, we assessed how interstitium stiffness altered expression of key glycocalyx glycoproteins. For these studies, adenocarcinomic human alveolar epithelial (A549) cells were grown on the platforms with either soft (∼5 kPa) or stiff (∼50 kPa) interstitium under either static or dynamic (cyclic 3-D stretch) conditions. As controls, A549 cells were also grown on tissue culture plastic (TC plastic). Once cells reached confluence, the A549 cells were collected, and the gene expression of key lung glycoproteins were assessed. Specifically, we examined the expression of the chondroitin sulfate proteoglycans versican (VCAN) and decorin (DCN) along with the heparan sulfate proteoglycans syndecans 1, 2, and 4 (SDC1, SDC2, and SDC4 respectively) using quantitative polymerase chain reactions (qPCR) (**Figure 2**).

**Figure 2:**
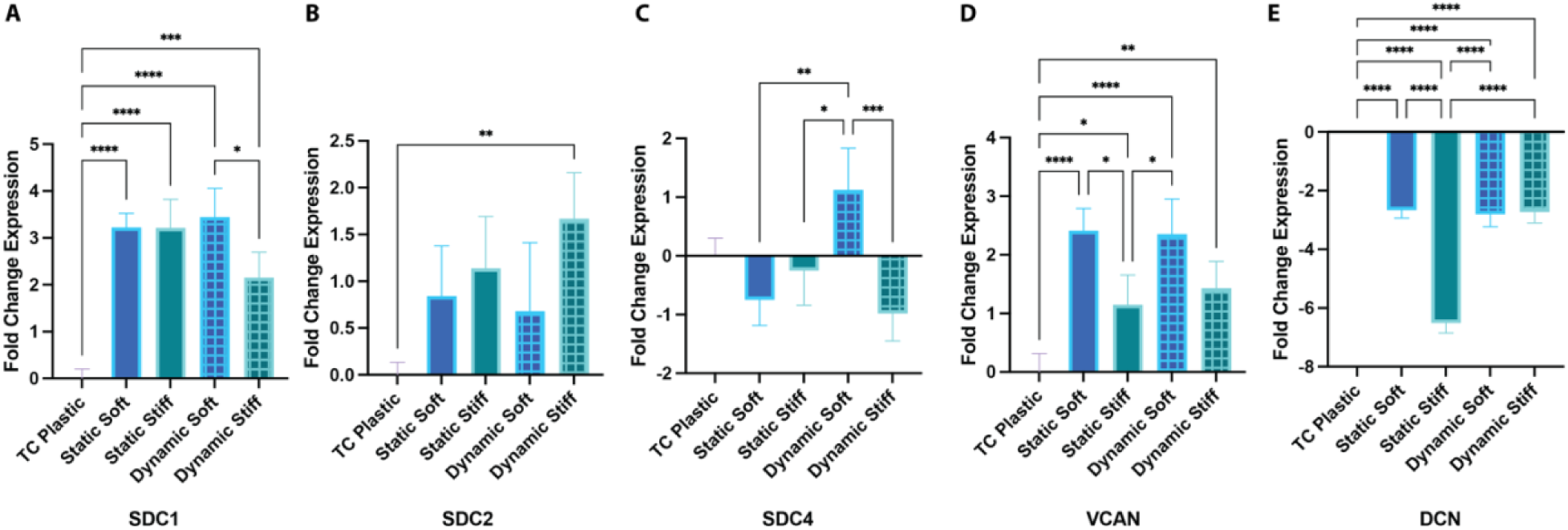
Effect of substrate properties on glycoprotein gene expression. **A-E)** Relative gene expression of SDC1 (**A**), SDC2 (**B**), SDC4 (**C**), VCAN (**D**), and DCN (**E**) in the epithelial cells grown on TC plastic, substrates with different stiffness that were under static or dynamic 3-D stretching conditions. All changes are reported relative to GAPDH expression and cells grown on TC plastic. **** p < 0.0001, *** p < 0.001, ** p < 0.01, and * p < 0.05; one-way ANOVA with Tukey HSD post-hoc analysis. All data is mean ± standard error of the mean.

qPCR analysis showed that biomimetic interstitium chips changed glycoprotein expression greatly in comparison to TC plastic. Specifically, SDC1 and VCAN were upregulated and DCN was downregulated compared to TC plastic when placed on our biomimetic platforms regardless of stiffness or whether dynamic stretch was applied (**Figure 2A, D-E**). SDC2 expression also increased in the cells grown on dynamic stiff substrates compared to TC plastic (**Figure 2B**). Together, these results indicate that TC plastic does not accurately reflect mechanically driven changes in glycoprotein genes.

In addition to changes relative to TC plastic, we observed significant changes in gene expression depending on the substrate stiffness and the presence of dynamic forces. Stiffness alone can affect glycoprotein gene expression. Specifically, elevated stiffness results in downregulation of VCAN and DCN compared to static soft conditions (**Figure 2D-E)**. However, stiffness related changes are not observed when dynamic forces are applied for these two genes. Specifically, we find that SDC1 and SDC4 expression is upregulated in cells grown on soft substrates compared stiff substrates when dynamic forces are applied, an effect that is not observed for static conditions. Conversely, increased expression of SDC4 is observed in cells on soft substrates only when dynamic forces are applied compared to other biomimetic substrates (**Figure 2C**). Similarly, stiffness dependent SDC1 gene expression changes are only observed when dynamic forces are applied (**Figure 2A**). Together, these results indicate that gene expression for glycoproteins are regulated by a complex combination of interstitium stiffness and dynamic forces. Neither stiffness nor dynamic forces are sufficient to reflect the diversity of gene expression changes observed here. To evaluate functional expression of glycoaminoglycans (GAGs), we performed an enzyme-linked immunosorbent assay (ELISA) on the glycocalyx collected from cells grown under different conditions (**Figure S1**). Our results show increased GAG expression, both for heparan sulfates and chondroitin sulfates, when cells are grown on biomimetic chips compared to TC plastic when pooled for all conditions. This is in agreement with our results at the proteoglycan level. Differences were also observed for chondroitin sulfate levels for static stiff on comparison to TC plastic and static soft conditions. However, there are no statistical differences for individual devices for heparan sulfates, most likely due to the sensitivity of the assay.

### Designing Peptide Functionalized Liposomes Targeting the Lunge Epithelium Glycocalyx

Based on our observations that glycocalyx expression changes as function of dynamic stretch and stiffness of the supporting substrate, we designed liposomal nanoparticles (LNPs) aimed at facilitating uptake into cells depending on the mechanical environment. Since we observed differences in the expression of different types of glycoproteins, we sought to create LNPs that recognize specific GAGs. Known peptide structures derived from viral sources that interact with chondroitin sulfate A (CSA), chondroitin sulfate C (CSC) or heparan sulfates (HS; see **Table 1** for sequences) were utilized for coating the LNPs. We note that these peptides (and related ones) often have cross-reactivity with other GAGs, but have chosen ones with higher affinity for each target.^61–63^ The peptides were chosen due to our observations that chondroitin and heparan sulfates are regulated by mechanical forces and substrate stiffness. For these studies, we modified the peptides to include cysteines to allow easy functionalization using bioconjugation strategies. To facilitate coating of the LNP surface, peptide-lipid conjugates were synthesized by reacting to a maleimide terminated polyethylene glycol (PEG 20 kDa) lipid with our cysteine terminated thiols in anhydrous conditions (**Figure S2**). Peptide conjugates were subsequently purified using HPLC. Synthesis of conjugates was confirmed by labelling the peptide-conjugates with fluorophores in a two-step process. First, amines on the peptides were labelled with methyltetrazine by incubating conjugates with a N-Hydroxysuccinimide ester (NHS) methyltetrazine. Second, a trans-cyclooctene fluorophore was reacted with peptides and the product were run in a polyacrylamide gel. As a control, lipids without peptides were treated in the same way and no conjugation was observed. These results indicate that we successfully synthesized the lipid-peptide conjugates (**Figure S3**).

**Table 1.** Peptides used for targeting with Primary Recognition GAGs

| Peptide Sequence | Primary Recognition GAG |
| --- | --- |
| CERRIWFPYRRF | Chondroitin sulfate C <sup>64</sup> |
| CRTPPESYASVR | Chondroitin sulfate A <sup>63</sup> |
| CGRKKRRQRRRPQ | Heparan sulfates <sup>65</sup> |

Next, we created base LNP structures consisting of 58 mol% 1,2-dioleoyl-sn-glycero-3-phosphocholine (DOPC) and 42 mol% cholesterol that were systematically extruded to 100-nm diameters. The lipid-peptide conjugates were then incubated with the pre-formed liposomes to allow for spontaneous intercalation into the liposome (**Figure 3A**). Dynamic Light Scattering (DLS) shows increased size upon incubation with peptide functionalized lipids (**Figure 3B**) that is consistent with the expected size shift for our peptide conjugates, indicating that we modified the liposome surface with the peptides.

**Figure 3.**
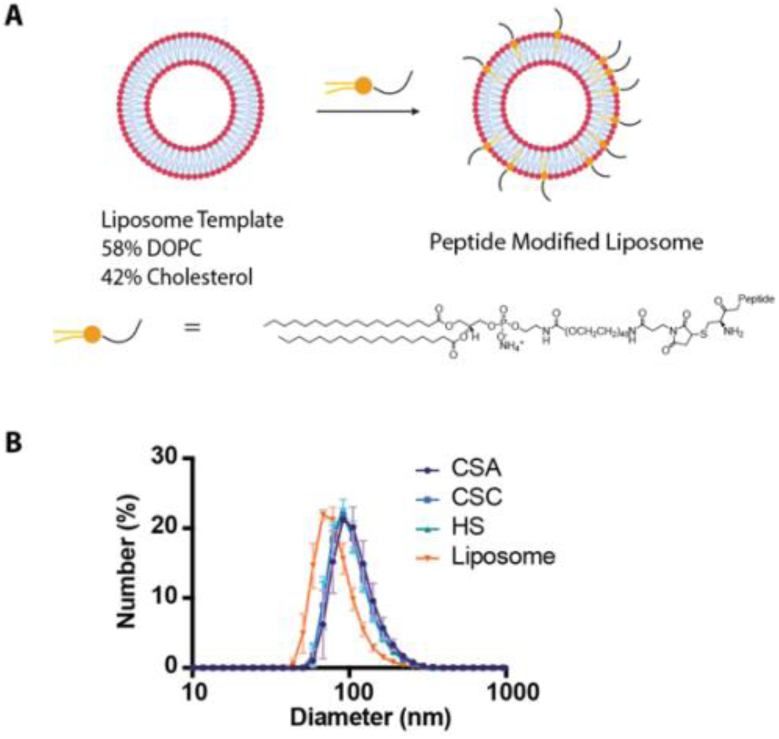
Design of nanoparticles for targeting glycocalyx changes. **A)** Schematic depicting the strategy for modifying liposomes with peptides that target different glycocalyx GAGs. **B)** Dynamic light scattering size distributions for bare liposomes (orange) along with peptides targeting chondroitin sulfate A (CSA, purple), chondroitin sulfate C (CSC, Blue), and heparan sulfate (HS, green).

### Effect of Mechanical Environment on Nanoparticle Uptake

Next, we assessed whether there were differences in LNP uptake depending on the surface peptide and the culture conditions. For these studies, the liposomes were formulated with lipophilic fluorophores. First, we assessed LNP uptake in conventional screening conditions (TC plastic). LNPs were incubated with A549 cells, and uptake was assessed via flow cytometry. In plastic conditions, unmodified liposome controls showed greater uptake than all targeting structures. HS LNPs enhanced uptake compared to the liposomes functionalized with CSC and CSA peptides (**Figure 4A**).

**Figure 4.**
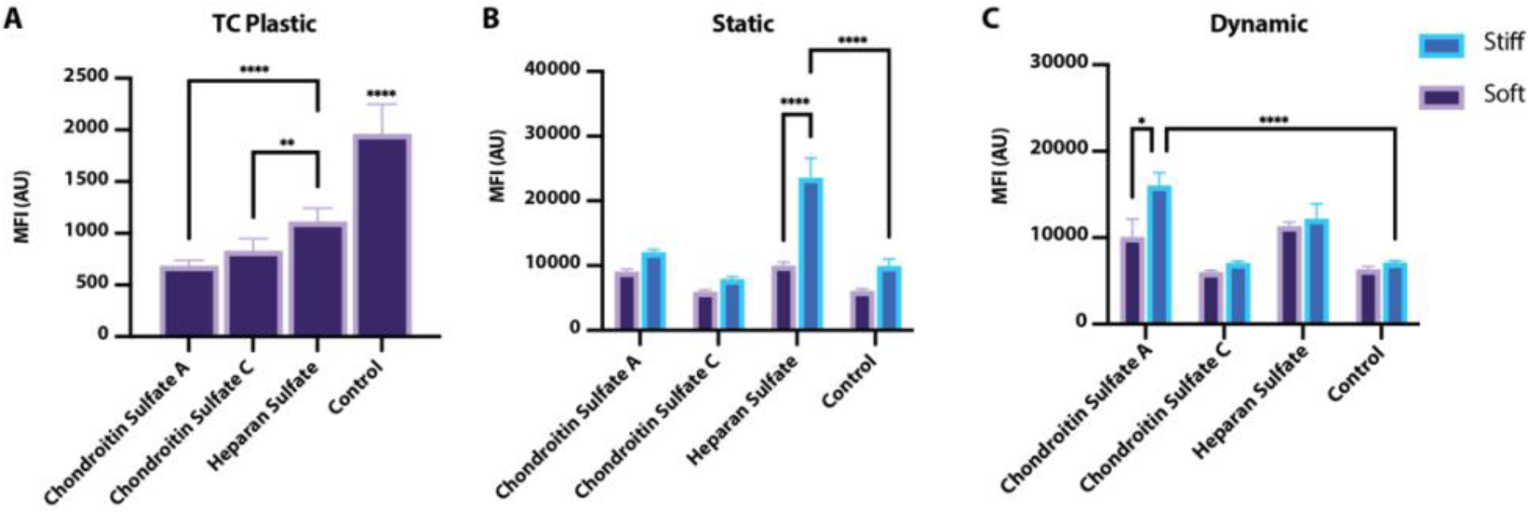
Uptake of LNPs into A549 cells in different mechanical environments. **A-C)** Flow cytometric median fluorescence intensities of A549 cells grown TC Plastic **(A)**, static soft or stiff substrates **(B)**, on soft or stiff substrates undergoing dynamic cyclic stretches **(C)** following treatment with fluorophore labelled LNPs formulated with different targeting peptides. **** p < 0.0001, *** p < 0.001, ** p < 0.01, and * p < 0.05; two-way ANOVA with Tukey HSD post-hoc analysis for comparing for substrate effects and relative to control LNP. All data is mean ± standard error of the mean.

To assess whether mechanical environment altered LNP uptake, A549 cells were cultured on the alveolar interstitium platforms for 1 week to allow the cells to reach monolayer confluency. Again, we utilized soft (5 kPa) and stiff (50 kPa) conditions with either dynamic forces or static cultures to assess how environment affected nanoparticle uptake. Under static conditions, only HS targeting LNPs showed greater uptake into A549 cells grown on stiff substrates in comparison to soft and control (non-targeting) liposome. They also showed significant increases in uptake compared to unmodified control LNPs (**Figure 4B**). In contrast, dynamic conditions showed no improvement in HS targeting LNPs for cells on soft or stiff substrates. There was also no improvement in comparison to control LNPs. When dynamic stretch was introduced to the A549 cells, CSA targeting LNPs showed enhanced uptake into cells grown on stiff substrates in comparison to the controls (unmodified liposomes; **Figure 4C**). These results conclusively show that mechanical stiffness changes the uptake preference of different LNP formulations and the addition of dynamic forces further alters uptake preference. In summary, HS targeting improves uptake into cells grown on stiff static substrates, while CSA targeting peptides enhances uptake into the cells grown on stiff substrates only when subjected to dynamic forces.

If we evaluate these results in the context of proteoglycan models, we observe that changes in proteoglycan content only partially explains nanoparticle uptake observation. We see that the upregulation of GAGs present in all our biomimetic models in comparison to conventional TC plastic can have a mixed effect. The lack of GAGs limits entry for targeted LNPs in all plastic conditions. However, increased GAG expression can either promote or inhibit uptake depending on the expression of surface GAGs cell surface. High levels of HS on cells grown in soft conditions do not necessarily improve entry for HS targeting LNPs. This result is in agreement with observations of others that GAG content is both necessary for entry with this peptide but potentially inhibitory at high levels.^65–68^ These observations, along with our results, point to the need to explore multiple combinations of targeting as we have here in order to identify high performing formulations.

## Conclusion

Alveolar epithelial cells are subjected to constant cyclic stretches and frequently encounter stiffer extracellular matrices (ECM) in a fibrotic or cancerous lung. We have devised a cell culture platform that mimics these dynamic and mechanical characteristics of ECM to study lung alveolar cells *in vitro*. Our studies demonstrate that the expression of cellular glycocalyx glycoproteins change depending on the mechanics and stretch state. Conventional culture conditions (tissue culture plastic) fail to accurately capture the glycocalyx changes driven by the physical environment. Our studies show that nanoparticle uptake is highly dependent on the physical environment. Whereby, screening of different formulations in biomimetic systems leads to the identification of different potentially therapeutically relevant LNP structures compared to conventional screening approaches. All these results together show the importance of considering ECM mechanics and dynamics when designing novel drugs/carriers. Future nanoparticle targeting studies *in vitro* would benefit from including physical attributes in targeting conditions. Collectively, it sets the stage for potentially bridging gaps between *in vitro* and *in vivo* models.

## Methods

### Safety Statement

No unexpected or unusually high safety hazards were encountered.

### Fabrication of Alveolar Interstitium

All the parts of the platform except the nanofibrous membrane were made by casting the mixture of PDMS (Sylgard 184, Dow Corning) resin and curing agent at a 10:1.05 ratio (w/w) on the 3-D printed molds, followed by a curing process at 75°C for 2 hours. Meanwhile, a thin PDMS membrane was prepared via spin-coating at 2500 rpm for a minute on a silicon wafer and curing at 75°C for an hour, and then bonded onto the pneumatic layer to construct the pneumatic chamber using a microtransfer assembly (µTA) technique that we previously developed.^52^ Briefly, the pneumatic layer with the opening facing down was stamped on a PDMS adhesive layer, which was prepared by spin-coating a PDMS mixture on a silicon wafer at 4000 rpm for 30 seconds, and transferred onto the PDMS membrane, followed by curing at 75 °C for 1 hour under a compressive pressure of approximately 1 MPa. The pneumatic layer with the bonded membrane was then gently peeled off from the silicon wafer. The middle interstitium layer was prepared by first bonding the interstitium layer onto the pneumatic layer with PDMS membrane using µTA technique and then adding Sylgard 527 (corresponding to 5 kPa normal lung tissue) or a 10:1 mixture of Sylgard 527 and 184 (50 kPa fibrotic tissue) into the interstitium chamber followed by curing at 75 °C for 4 hours.

Prior to cell culture, the electrospun nanofibrous membrane was bonded to the cell culture layer using the µTA technique, treated with oxygen plasma at medium power setting for 1 min in a plasma cleaner (Model PDC-001, Harrick Plasma), and bonded onto the interstitium layer. At last. Subsequently. phosphate buffered saline (PBS) (BioWhittaker, Lonza) was added to the cell culture chamber to keep the fibrous membrane immersed and the whole platform was cured at 45 °C overnight.

### Electrospinning and Characterization of Nanofibrous Membranes

A solution of PCL in 1,1,1,3,3,3-hexafluoro-2-propanol (HFIP, 10%, w/v) was electrospun to produce the nanofibrous membrane. Briefly, the PCL solution was loaded into a syringe with a gauge blunt-tipped needle as the spinneret and ejected using a syringe pump (New Era Pump Systems Inc) at a flow rate of 0.5 mL/hour. A square glass plate (4” x 4”) was placed on an aluminum tap covered plate served as the fiber collector, which was displaced 16 cm away from the spinneret and grounded to the power supply. A voltage of 14 kV was applied to the spinneret using a DC power supply (Spellman Bertan) for 20 min.

The nanofibrous membrane was sputter-coated by gold (Denton Vacuum LLC) and images were taken using a scanning electron microscope (SEM, TM3030 Plus, Hitachi High-Technologies Co., Tokyo, Japan). The pore size and fiber diameter were analyzed using ImageJ. Briefly, the images were adjusted using threshold command. The area of each pore was analyzed using the function of analyze particles. The diameter of fibers was measured by drawing a line across the fiber and measuring the length. Then, a histogram line graph was generated using Prism 8 (GraphPad software) to show the distribution of the measured pore areas or fiber diameter.

### Characterization of Mechanical Stretching

The PDMS membrane bonded on the pneumatic chamber or the platform without the cell culture chamber were used for the mechanical stretch characterization. When air was infused into and withdrawn from the pneumatic chamber, the deformation (strain) of the PDMS membrane with/without the nanofibrous membrane immobilized on the PDMS interstitium was characterized using the same method we reported previously.^51^ Briefly, the deformation of the membranes was captured using a charge-coupled device (CCD) camera (Model DMK 31, The Imaging Source) and the curvilinear profiles of the membranes were obtained using ImageJ (http://rsb.info.nih.gov/ij/index.html). The theoretical calculation of the linear strain was also detailed previously.^51^

### Cell Growth

Human alveolar epithelial cells (A549; Cat#: CCL-185, ATCC) were cultured in Dulbecco’s Modified Eagle Medium (DMEM) with L-glutamine (Life Technologies) supplemented with 10% fetal bovine serum (FBS), 100 U/ml penicillin and 100 μg/ml streptomycin (Life Technologies). The nanofibrous membrane was coated with 50 μg/ml type I collagen (Col I, rat tail, Corning) overnight in the incubator. The cells were seeded at a density of 1 × 10^5^ cells/cm^2^ and cultured in the incubator of 37 °C and 5% CO_2_. The cells were cultured under static condition for 1 day, and then subjected to mechanical stretch continuously for 5 days and culture medium were refreshed every other day.

### qPCR Study of Cells Cultured in Different Conditions

To observe cellular glycocalyx expression changes in static and dynamic culture interstitiums qPCR was performed. RNAqueous Micro Kit (Invitrogen by Thermo Fisher Scientific) was used for RNA isolation following standard working protocol. Briefly, RNA isolation was confirmed in NanoDrop one (Thermo Scientific). RNA samples were mixed with SuperScript IV VILO Master Mix with ezDNase (Invitrogen by Thermo Fisher Scientific) kit solutions to make cDNA library. Using SimpliAmp Thermal Cycler (Applied Biosystems by thermo fisher Scientific), cDNA library was prepared. The primers for SDC1, SDC2, SDC4, VCAN, DCN and GAPDH (TaqMan Gene Expression Assays, Applied Biosystems by Thermo Fisher Scientific) expression and TaqMan Fast Advanced Master Mix (Applied Biosystems by Thermo Fisher Scientific) was used with cDNA samples for qPCR. Standard protocols were followed for mixing, plating in a 384 well plate, and running qPCR on a LightCycler 480 (Roche).

### Enzyme Linked Immunosorbent Assay (ELISA) Study of Cells Cultured in Different Conditions

Cells cultured in TCP, static and dynamic culture interstitiums were dissociated using 0.25% trypsin (with 2.21mM EDTA, 1X [-] sodium bicarbonate) solution (Corning). Then they were collected and mixed with previously formulated cell culture medium (DMEM with L-glutamine supplemented with FBS, penicillin and streptomycin) (see methods for cell growth), maintaining 1:1 volume ratio with trypsin solution. The mixture was then centrifuged at 500xg for 5 mins and cell supernatant was collected. The supernatant was used as samples for CS ELISA kit (TSZ Scientific, Biotechnology systems) and HS ELISA kit (TSZ Scientific, Biotechnology systems). Standard working protocol of the kits was followed for the assay. Cytation 5 imaging reader (BioTek) was used to read the plate for absorbance at 450nm when the assay was complete.

### Peptide-Lipid Conjugate Synthesis

Peptides, purchased from Genscript with >98% purity, terminated with cysteines were reacted with twice the molar amount of 1, 2-Distearoyl-sn-glycero-3-phosphoethanolamine-Poly (Ethylene glycol) (DSPE-PEG20kD) maleimide in anhydrous DMSO with triethylamine overnight at 4 °C. Samples were purified on an Accela ultra-high performance liquid chromatographer with a gradient of triethylammonium acetate (TEAA) buffer (0.05 M; pH 7.0) to 95% acetonitrile with TEAA buffer (0.05 M; pH 7.0) on a biobasic C18 column (Thermo Electron Corporation). The concentration of peptide conjugates was determined by the standard Bicinchoninic acid (BCA) assay using Pierce(tm) BCA Protein Assay Kit (Thermo Fisher Scientific). To confirm successful synthesis of peptide conjugates, they were reacted with 10:1 Methyltetrazine-sulfo-N-Hydroxysuccinimide ester (Met-NHS) (Click Chemistry Tools):Peptides for an hour at room temperature in Dulbecco’s Phosphate Buffered Saline (DPBS) (BioWhittaker, Lonza). The conjugates were further incubated with 1.1:1 Cyanine 5 trans-cyclooctene (AAT Bioquest):Met-NHS for a minute and run through NuPAGE 4-12% Bis-Tris premade polyacrylamide gel (Invitrogen by Thermo Fisher Scientific). The mini gel tank (Invitrogen by Thermo Fisher Scientific) and PowerEase 300W (Life Technologies) system was used for running the electrophoresis. Electrophoresis ran for 1 hour at 120V to ensure complete running of load through the gel. The Gel was imaged in ChemiDoc MP Gel Imaging System (BioRad Labnoratories Inc).

### Liposome Synthesis

Liposomes were synthesized following the widely used thin film technique with 58 mol% 1,2-dioleoyl-sn-glycero-3-phosphocholine (DOPC; Avanti Polar Lipids) and 42 mol% cholesterol (Sigma Aldrich). Liposomes were synthesized by forming lipid cakes by removing chloroform under vacuum for >30 min followed by 2 h on a 4.5 L, -105 °C lyophilizer (Labconco). Lipids were reconstituted in PBS and sonicated for 20 minutes. Liposomes were systematically sized to by extruding through a 100 nm membrane for 15-20 passes (Nuclepore Track-Etch Membrane, Cytiva). For flow cytometry studies, the lipid formulations were synthesized to include 0.1 mol% of either Vybrant DiI or DiO (Invitrogen), a highly lipophilic dye.

### Peptide Decorated Liposome Preparation

300 peptide-lipid conjugates were added for one liposomal vesicle and incubated overnight at 4 °C. Dynamic Light Scattering (DLS) was performed on liposomes with a Zetasizer S (Malvern Instruments).

### LNP Uptake of Cancerous Lung Epithelium Cells

Lung Epithelium cell line A549 were incubated with 0.15 nM Chondroitin sulphate A, Chondroitin sulphate C and Heparan sulphate decorated and non-decorated liposomes for 4 hours in DMEM. The incubation was initiated after reaching monolayer confluency of cells on their substrates. Uptake efficiency was observed for soft, stiff matrices in both static and dynamic conditions and compared with plastic control.

### Flow Cytometry of LNP uptake

To quantify the accumulation of the peptide decorated liposomes inside the cells, flow cytometry was performed after 4-hour incubation of the cells with liposomes. Cells were collected by trypsonization after 4 hours, rinsed 3 times with 1x Dulbecco’s phosphate buffered saline (DPBS) (Formulation: 200mg/L KCl and KH_2_PO_4_, 8000mg/L NaCl, 2160mg/L Na_2_HPO_4_*7H_2_O, pH 7-7.6,) (BioWhittaker, Lonza) fixed with 3.7% paraformaldehyde solution prior to running in flow cytometer (Guava easyCyte 6HT, Luminex). The fixing solution was prepared by mixing 10% formaldehyde (36.5-38% Formaldehyde solution in H_2_O, Sigma) and 89% DPBS. 1% 1.5M NaCl solution prepared with Sodium Chloride crystals (Fisher Bioreagents) in H_2_O was added to balance the pH level of the solution.

### Statistical Analysis

All statistical analysis was performed with GraphPad PRISM 9 (GraphPad Software, Inc.). All data sets were assessed with Welch’s t-test (Figure S1a,c), one-way ANOVA followed by a Tukey HSD *post hoc* analysis (Figures 2, 4a, and S1b,c) or a two-way ANOVA followed by a Tukey HSD *post hoc* analysis (Figure 4b,c). Significance thresholds were *α=0*.*05*. Statistical parameters are as follows: Figure 2: (A: F = 33.27, B: F = 5.37, C: F = 9.49, D: F = 18.49. E: F = 198); Figure 4 (A: F = 95.87 B: F_interaction_ = 10.3, F_column_ =44.08, F_row_ = 28.04; C: F_interaction_ = 2.55, F_column_ =7.4, F_row_ = 18.53); Figure S1 (B: F = 1.56, D: F = 5.46).

## Author Contributions

The manuscript was written through contributions of all authors. All authors have given approval to the final version of the manuscript.

## Notes

The authors declare no competing financial interest

## Acknowledgements

Research reported in this publication was supported by the Oak Ridge Associated Universities (B.M.), University of North Texas Seed Funds (B.M. and Y.Y), and the National Institutes of Health under awards R21GM141563 (B.M.) and R15EY033967 (Y.Y).

TOC: The effect of substrate stiffness and dynamic cyclic stretch on lung epithelial glycocalyx expression was examined using biomimetic cell culture techniques. Nanoparticles were designed to target glycocalyx changes, and their uptake by epithelial cells in different physical environments was evaluated.

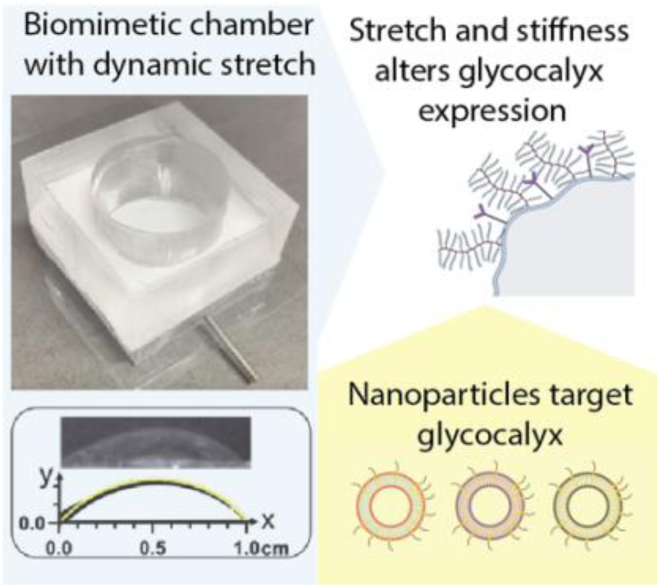

## Notes

### Competing Interest Statement

The authors have declared no competing interest.

