## Supplemental Files for "Nanoparticle targeting of mechanically modulated glycocalyx"

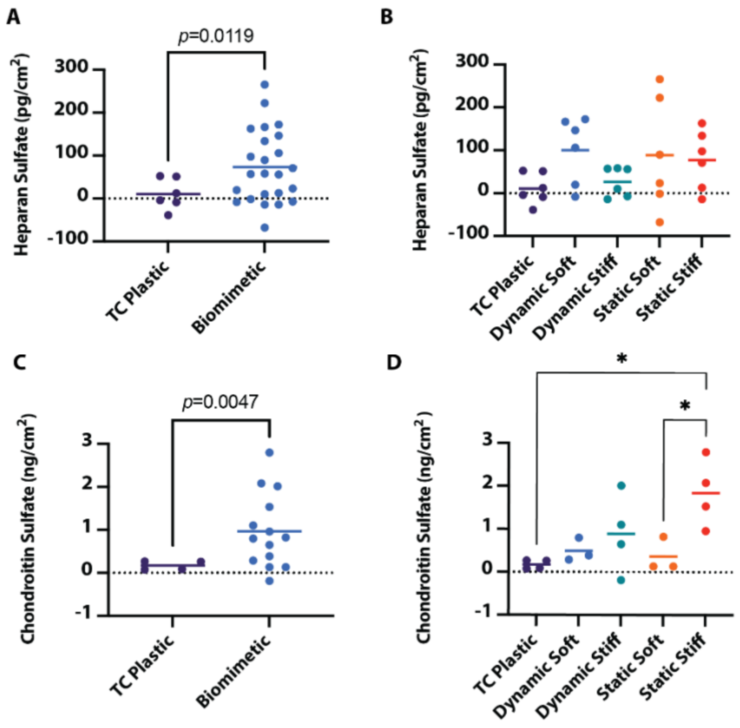


**Figure S1. GAG content in the glycocalyx measured through ELISA. A)** Heparan sulfate content in the cell glycocalyx grown on TC plastic in comparison to biomimetic chip samples. **B)** Heparan sulfate content in the cell glycocalyx for different device types. **C)** Chondroitin sulfate content in the cell glycocalyx grown on TC plastic in comparison to biomimetic chip samples. **D)** Chondroitin content in the cell glycocalyx for device types. A,C: Welch's t-test; B,D: one-way ANOVA with Tukey HSD *post hoc*; * $p < 0.05$. Bars show mean.

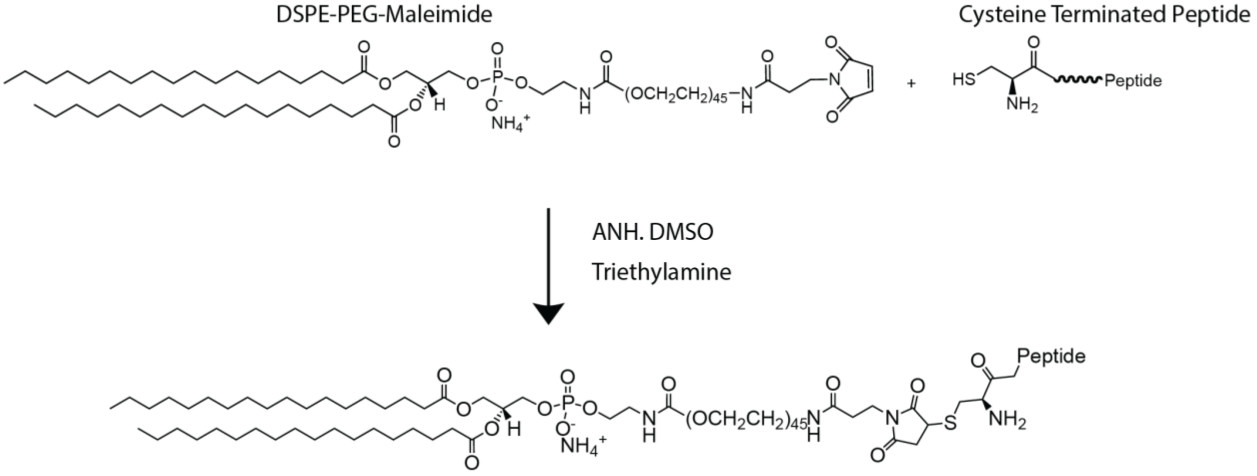


**Figure S2. Schematic depicting peptide conjugation to lipids.** Cysteine terminated peptides are reacted with DSPE-PEG-Maleimide in anhydrous DMSO overnight.

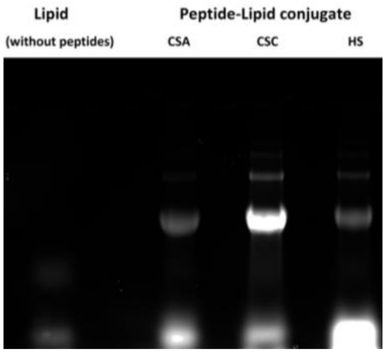


**Figure S3. Electrophoresis of glycocalyx targeting peptide-lipid conjugates in comparison to non-conjugated lipid control.** Polyacrylamide gel electrophoresis of CSA, CSC and HS targeting peptide-lipid conjugates after reacting with N-Hydroxy succinimide ester (NHS) methyl tetrazine and transcycloctyne fluorophore confirmed the presence of peptide lipid conjugate bands. No conjugation band was observed after a similar reaction with lipids.
